# Tightly-orchestrated rearrangements govern catalytic center assembly of the ribosome

**DOI:** 10.1101/444414

**Authors:** Yi Zhou, Sharmishtha Musalgaonkar, Arlen W. Johnson, David W. Taylor

## Abstract

The catalytic activity of the ribosome is mediated by RNA, yet proteins are essential for the function of the peptidyl transferase center (PTC). In eukaryotes, final assembly of the PTC occurs in the cytoplasm by insertion of the ribosomal protein Rpl10. We determine structures of six intermediates in late nuclear and cytoplasmic maturation of the large subunit that reveal a tightly-choreographed sequence of protein and RNA rearrangements controlling the insertion of Rpl10. We also determine the structure of the biogenesis factor Yvh1 and show how it promotes assembly of the P stalk, a critical element for recruitment of GTPases that drive translation. Together, our structures provide a blueprint for final assembly of a functional ribosome.

**One Sentence Summary:** Cryo-EM structures of six novel intermediates in the assembly of the large ribosomal subunit reveal mechanism of creating the catalytic center.

## Main Text

Ribosomes are the molecular machines that all cells depend on for protein synthesis. Its two fundamental functions, decoding and peptide synthesis, are separated into the small and large ribosomal subunits, respectively. Despite using RNA for catalysis, ribosomes are ribonucleoprotein particles, and proteins surrounding the peptidyl transferase center (PTC) are essential for function. In eukaryotes, ribosomal subunits are largely preassembled in the nucleus (*1-5*) but exported in a functionally inactive state to the cytoplasm (*6-9*), where final assembly of the large (60S) subunit follows a hierarchical pathway (*10*). Two critical events of 60S maturation are completion of the PTC by insertion of Rpl10, which primes the subunit for a quality control “test drive” (*11, 12*), and assembly of the P stalk that recruits and activates the GTPases of the translation cycle (*13*). While extensive molecular genetics and biochemical studies over the past 20 years have revealed clues into this complex process, the mechanisms for assembly of both the PTC and the P stalk have remained elusive. Here, we have determined the structures of a series of intermediates of 60S maturation that reveal the dynamic changes in RNA conformations and protein exchanges required for assembly of the P stalk and completion of the PTC.

### Prying open of RNA helices H38 and H89 primes the 60S for Rpl10 loading

Using cryo-electron microscopy (cryo-EM) of 60S precursors trapped from yeast, we captured six intermediates in late nuclear and cytoplasmic 60S assembly at ~3.5-3.8 Å resolution (Fig.1, figs. S1 to S4, Movie S1 and S2), which provide unprecedented mechanistic insights into 60S maturation. To isolate particles immediately after export from the nucleus, we used tandem affinity purification (TAP)-tagged mutant Rlp24 (Rlp24∆C), which fails to recruit the AAA-ATPase Drg1 that initiates cytoplasmic maturation (*14*). Surprisingly, we identified both late nuclear (LN) and early-cytoplasmic (EC) particles, the latest nuclear and earliest cytoplasmic particles visualized to date, respectively. The pre-export LN particle lacks the nuclear export adapter Nmd3 but contains the biogenesis factors Rlp24, Bud20, Mrt4, Nog1, Nsa2, Nog2, Tif6 and Arx1 (Fig. 1A). We also observe clear density for Rpl12 on the P stalk. The L1 stalk, which interacts with E site ligands in a mature ribosome, is in an “open” conformation and H89, which constitutes part of the binding cleft for Rpl10, is displaced by the N-terminal domain (NTD) of Nog1 in the A site. In addition, H69, which provides a critical intersubunit bridge in the complete 80S ribosome, is disrupted by Nog2. Earlier studies of other nuclear pre-60S intermediates showed that in both the Nog2 and Arx1 particles (*15, 16*), the 5S has not yet rotated into its mature position and internal transcribed spacer 2 (ITS2) has not been processed. In the LN structure, the 5S RNA has rotated into its mature position and ITS2 has been removed, placing the LN particle later than the Arx1 and Nog2 particles in maturation. The presence of Rpl42, which completes the binding site for Nmd3 (*17, 18*), and Rpl29 in the LN particle also place it later in the assembly pathway than the previously described Rix1 particle (*19*). Thus, the LN particle represents a pre-60S particle immediately before gaining export competence by exchanging Nog2 with Nmd3 (*20*) to drive export to the cytoplasm.

**Fig. 1.**
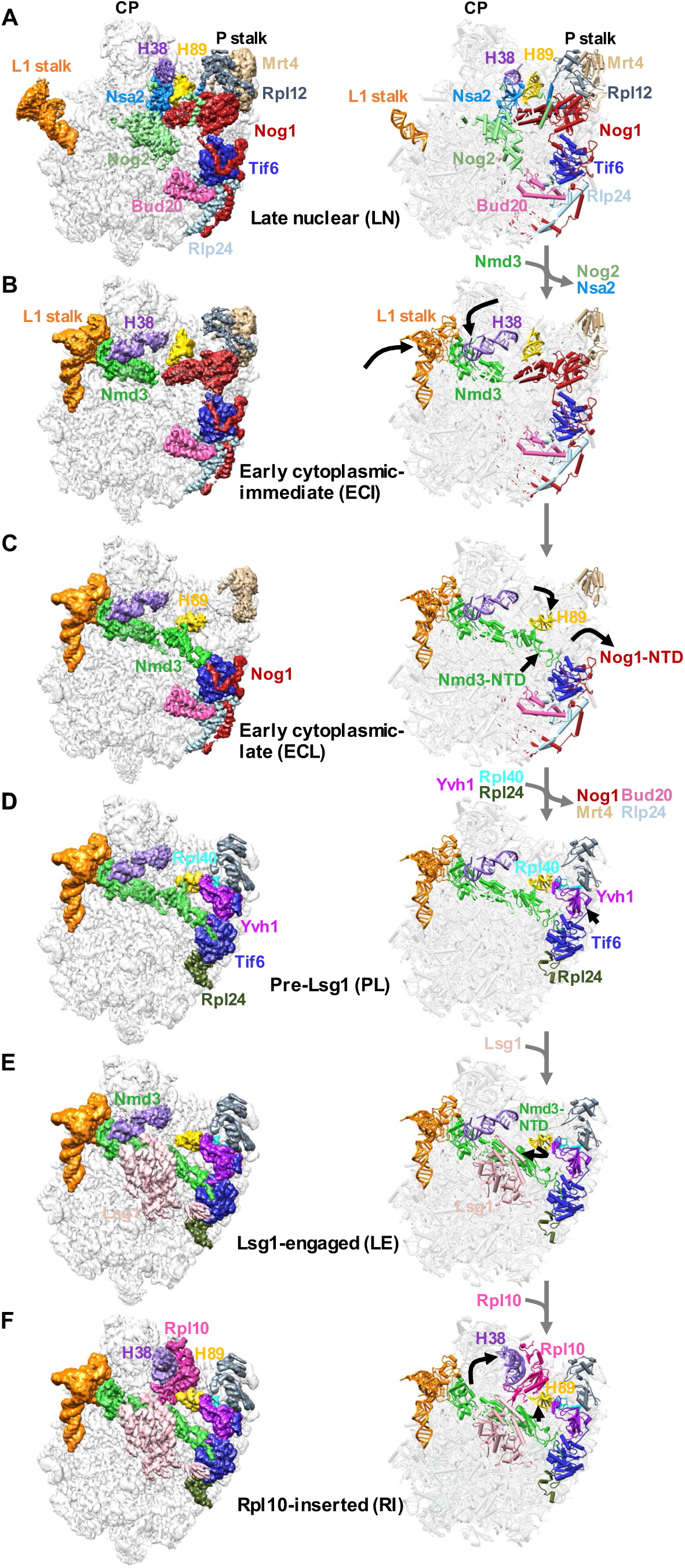
Structures of late nuclear and cytoplasmic particles reveal the pathway of 60S maturation. Crown view of late nuclear (A), early cytoplasmic-immediate (B), early cytoplasmic-late (C), pre-Lsg1 (D), Lsg1-engaged (E), and Rpl10-inserted (F) ribosomal intermediates at ~3.5–3.8 Å resolution, with corresponding atomic models shown on the right. Key protein and rRNA elements are colored as indicated. Black arrows indicate conformational changes while grey arrows indicated protein exchanges. CP, central protuberance. In the ECI particle, Rpl12 was not modeled due to poor density. In the ECL particle, Rpl12 is not clearly discernible and Mrt4 is poorly resolved due to the mobility of the P stalk after the NTD of Nog1 is released from the A site. Therefore, Rpl12 was not modeled while Mrt4 was only partially modeled.

In the progression from the LN to EC particles, Nmd3 has exchanged with Nog2 for export from the nucleus and Nsa2 has been released (Fig. 1B). Nmd3 binding in the E and P sites promotes two large-scale rearrangements of RNA: closure of the L1 stalk and capture of H38, which constitutes the second RNA helix of the Rpl10 binding cleft. The L1 stalk is held closed by interactions between Rpl1 and the eIF5A-like domain of Nmd3 in the E site. H38 fits snugly into a saddle-shaped surface of Nmd3 where it is stabilized by multiple contacts including Arg333 stacking on the flipped out A1025 extending from the tip of H38. Strikingly, in this state, the tip of H38 is shifted ~50 Å from its position in the mature subunit (fig. S5A). Because H38 provides extensive contacts for Rpl10 in the mature subunit, this bending of H38 by Nmd3 partially opens the Rpl10 binding site to initiate priming of the pre-60S particle for loading of Rpl10.

The EC particles could be further separated into two states: early cytoplasmic-immediate (ECI) and early cytoplasmic-late (ECL). In the ECI particle, the NTD of Nog1 remains inserted in the A site, displacing H89 (fig. S5B) and preventing Rpl10 insertion into this particle. In addition, the ECI particle is the earliest particle in which H69 is folded into its nearly-mature conformation. As this coincides with Nmd3 binding, Nmd3 may promote H69 folding. However, Nmd3 also appears to drive the flipping of two bases, G2261 and U2269, (fig. S5C), distinguishing this structure from mature H69 and priming the particle for activation of the GTPase Lsg1, the release factor for Nmd3 (*18, 21*). Interestingly, in the ECI particle, there is no discernible density for the NTD of Nmd3 (Fig. 1B), most likely because it is highly mobile before docking onto Tif6.

In the ECL particle the N-terminus of Nog1 has been displaced from the A site, allowing the NTD of Nmd3 to dock onto Tif6 (Fig. 1C). H89 now rearranges into its nearly-mature position where it engages with a small loop within the NTD of Nmd3, which we name the histidine thumb (fig. S5B and Movie S3). Comparing the ECI and ECL particles shows that release of Nog1 is necessary to allow Nmd3 docking (fig. S5D). Surprisingly, the C-terminus of Nog1 remains in place on the ECL particle (Fig. 1C), which may inhibit the binding of Lsg1. Together, the interaction of the histidine thumb of Nmd3 with H89 and the capture of H38 by the eIF5A-like domain of Nmd3 hold open the Rpl10 binding site, priming the pre-60S for the loading of Rpl10.

To capture a series of particles downstream of the ECL particle, we partially arrested ribosome biogenesis *in vivo* with diazaborine, an inhibitor of the AAA-ATPase activity of Drg1 (*22*). The three classes of particles we obtained differ in their occupancy of Rpl10 and Lsg1 and show a large-scale structural rearrangement in Nmd3. In all three classes, Nog1, Bud20, Mrt4 and Rlp24 have been released while Rpl40 and Rpl24 have loaded. The pre-Lsg1 (PL) particle contains Nmd3 with the NTD interacting with H89 similarly to the ECL particle; the Lsg1 engaged (LE) particle contains Nmd3 and Lsg1 with the NTD of Nmd3 rotated to engage Lsg1; and the Rpl10-inserted (RI) particle contains Nmd3, Lsg1 and Rpl10 with the NTD of Nmd3 remaining engaged with Lsg1 (Fig. 1, D to F). Intriguingly, in all classes we observed a well-defined density between Tif6 and the P stalk (fig. S6A) that we identified as the zinc-binding domain (ZBD) of Yvh1. Yvh1 has 364 amino acids, comprising an N-terminal phosphatase domain (aa1-214) and a C-terminal ZBD (aa215-364). While the function of the phosphatase domain is unknown, the ZBD promotes Mrt4 release for P stalk assembly (*23, 24*).

### Yvh1 releases Mrt4 by rearranging the P stalk

We were able to build an atomic model of the ZBD of Yvh1 *ab initio* (fig. S6). The ZBD of Yvh1 is wedged between the P stalk and Tif6 and centered on the sarcin-ricin loop (SRL), an RNA element that is essential for activating translational GTPases. An extended internal loop of Yvh1 also engages the tip of H89 (Fig. 2A). The ZBD is composed almost entirely of beta-sheets and contains two predicted zinc-binding centers (fig. S6, C to E). Yvh1 interacts directly with the P stalk, H89 and the SRL, with Trp329 stacking on G1242 of H43, R269 stacking on C2284 of H89 and Phe260 stacking on A3027 of SRL (Fig. 2, B to D). An additional density adjacent to the P stalk that may account for the N-terminal phosphatase domain of Yvh1 can be seen at lower thresholds (fig. S7A). The C-terminus of Yvh1 was less well resolved but could be traced to the surface of Tif6 at lower thresholds (fig. S7B), where it interacts with the extreme N-terminus of Nmd3. Interestingly, the C-terminal 20 amino-acid tail of Tif6 folds back over itself to interact with Nog1 in the Nog1-bound particle, but reorients toward Yvh1 in the Yvh1-bound particle (fig. S7C). Comparison of the Nog1-containing ECI particle with the Yvh1-bound PL particle reveals that the NTD of Nog1 (fig. S7D) and the displacement of H89 by Nog1 (fig. S5B) would both occlude Yvh1 binding. Thus, Nog1 controls the timing of Yvh1 binding after export to the cytoplasm.

**Fig. 2.**
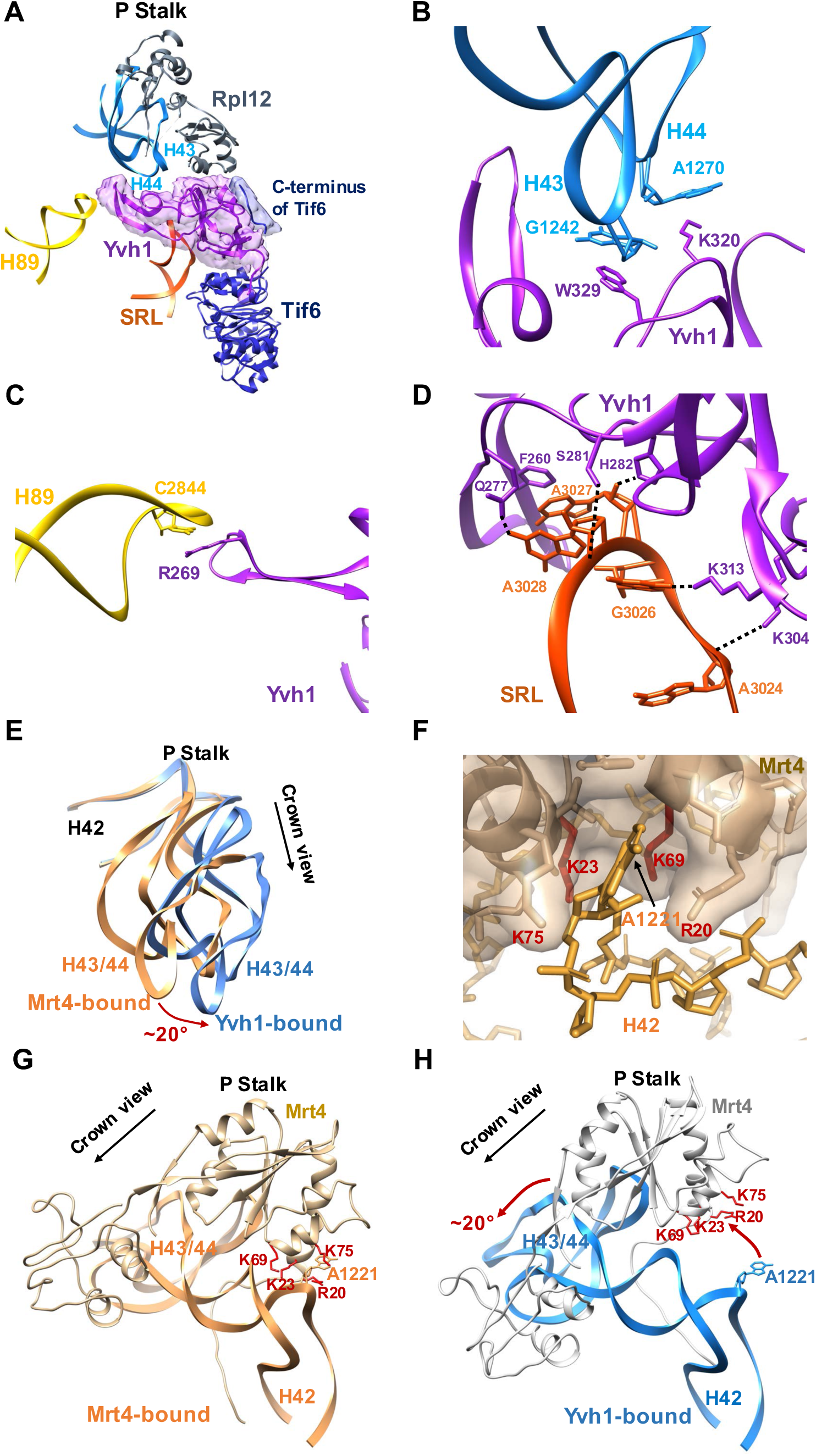
Yvh1 forces the release of Mrt4. **(A)** Atomic structure of the zinc binding domain of Yvh1 modeled into the unassigned density. The small piece of extra density (blue) was assigned to the C-terminus of Tif6 (see fig. S7B). **(B-D)** Interactions of Yvh1 with helices H43 and H44 of (B), helix H89 (C), and SRL (D). Dashed lines, predicted hydrogen bonds. **(E)** Comparison of the P stalk RNAs in the Mrt4-bound (orange) and Yvh1-bound (blue) states. **(F)** Mrt4 in cartoon and transparent surface representation. A1221 of H42 binds in a pocket of Mrt4. **(G)** Mrt4 bound to P stalk RNA rotated ~120° clockwise relative to the view in (A) showing A1221 of H42 in the binding pocket of Mrt4. Residues of Mrt4 interacting with H42 are indicated. **(H)** Mrt4 from the ECI particle was docked onto H43 and H44 of the Yvh1-bound PL particle, showing the movement of Mrt4 away from A1221 of H42.

To understand how Yvh1 releases Mrt4 for P stalk assembly, we compared the P stalk from Mrt4-containing ECI particles and Yvh1-containing PL particles. Intriguingly, we find that Yvh1 binding leads to allosteric changes in the binding site of Mrt4 to release this protein. The LN particle along with the ECI and ECL particles contain Mrt4 on the P stalk, whereas the PL particle lacks Mrt4 but contains Yvh1, consistent with previous work showing that the binding of Yvh1 displaces Mrt4 (*23, 24*). Notably, in the PL particle, the P stalk has undergone a rotation of ~20 degrees away from the Central Protuberance toward Tif6 (Fig. 2E). This rotation involves a rigid body rotation of the tip of the stalk (H43 and 44) with Rpl12 that is accomplished by bending of H42, which forms the stem of the stalk. The docking site of Mrt4 includes H43 and H44 as well as A1221 of H42, which inserts into a pocket in Mrt4 (Fig. 2F). Aligning H43, H44 and Rpl12 from the Mrt4-bound and Yvh1-bound structures, reveals that the Yvh1-induced bending of the P stalk RNA retracts the Mrt4 pocket away from A1221 on H42 (Fig. 2G and H), thereby reducing Mrt4 contacts with RNA. Mutations in Mrt4 that reduce its affinity for RNA bypass the need for Yvh1 (*23, 24*). These mutations include residues K23 and K69 that interact with A1221 (Fig. 2F), demonstrating that the interaction of Mrt4 with this RNA element is critical for its stable binding to the P stalk. Thus, Yvh1 controls the release of Mrt4 by allosterically altering its binding site, allowing final assembly of the P stalk proteins and setting up the subunit for a GTPase-dependent test drive.

### A large-scale rotation of Nmd3 releases H89 for Rpl10 loading

In the transition from the ECL particle to the pre-Lsg1 (PL) particle, the NTD of Nmd3 remains docked on Tif6 (Fig. 1D). The NTD of Nmd3 contains two zinc-binding centers (Fig. 3A). Residues 16-39 form a compact treble-clef that binds to Tif6 with the N-terminal residues of Nmd3 extending over the surface of Tif6. The C-terminal end of this zinc center contains a short alpha helix that connects to the remainder of the N-terminal domain through a single loop of amino acids. This is mirrored on the distal end of the NTD, where the second zinc-binding treble clef, comprised of C56-C58-C143-C145, connects to the eL22-like domain of Nmd3 in the P site through another loop of amino acids. Thus, the bulk of the NTD of Nmd3 is suspended by loops on each end, allowing this domain to pivot.

**Fig. 3.**
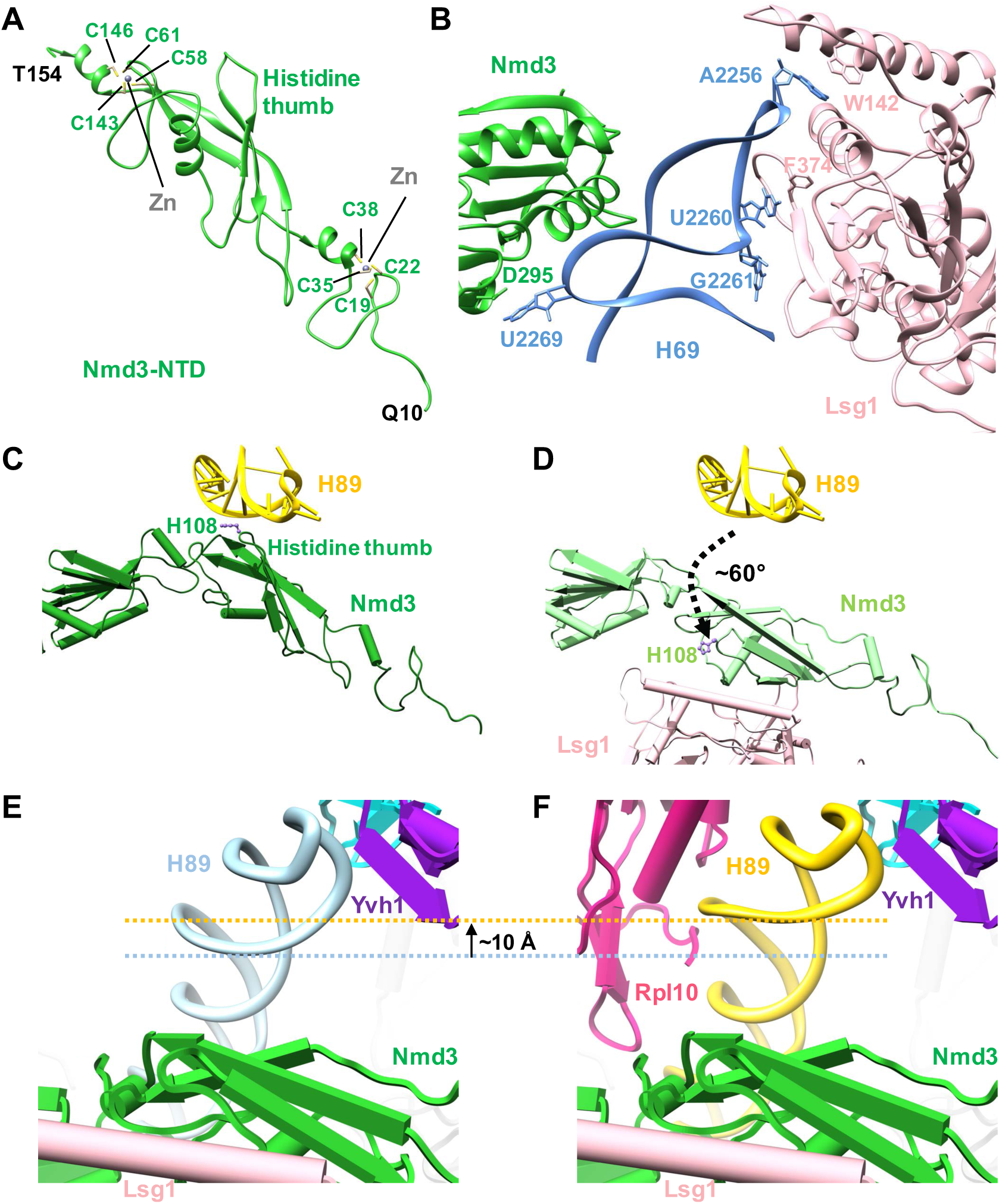
Large-scale rearrangements of Nmd3 NTD primes the subunit for Rpl10 insertion and completion of the catalytic center. **(A)** Atomic structure of the zinc-binding N-terminal domain of Nmd3. Cysteines of the two treble clef zinc binding motifs are indicated. Zn^2+^ ions were modeled into their predicted positions. **(B)** Interactions of H69 with Nmd3 and Lsg1 showing flipped out G2261 and U2269. **(C)** In the pre-Lsg1 particle, H108 of the Nmd3 histidine thumb is engaged with H89. **(D)** Upon Lsg1 binding, the bulk of the N-terminal domain of Nmd3 rotates ~60°, releasing the histidine thumb from H89. **(E-F)** Upon Rpl10 insertion, the middle portion of H89 is retracted ~10 Å to accommodate and stabilize Rpl10 in its binding cleft; the tip of H89 remains in position to interact with Yvh1.

In the Lsg1-engaged (LE) particle, Lsg1 interacts extensively with H69 with Trp142 stacking on A2256 at the tip of the helix and Phe374 stacking on U2260. The two flipped bases of H69, G2261 and U2269, interact with a strand of Lsg1 adjacent to Switch 1 of its GTPase center and Asp295 of Nmd3, respectively (Fig. 3B). Remarkably, the NTD of Nmd3 undergoes a ~60-degree rotation upon Lsg1 binding (Fig. 1E and Fig.3, C and D). In this rotated state, the histidine thumb has swung away from H89 to engage with the long alpha helix (residues 149-173) of Lsg1. This rotation involves swiveling of the N-terminal domain about the two linkers that connect this domain on one end to the zinc center, docked on Tif6, and on the other end, to the eL22-like domain in the P site. This rotation is also accompanied by a slight displacement of the C-terminal end of the zinc-binding domain of Nmd3 toward Lsg1 (Movie S4). Importantly, the release of H89 from the histidine thumb of Nmd3, frees this helix to adopt its final, mature position upon Rpl10 insertion. Therefore, the LE particle is “primed” for Rpl10 loading by both the prying open of H38 and release of H89 by Nmd3.

### Release of H38 and H89 from Nmd3 stabilizes Rpl10 in the ribosome

Ultimately, insertion of Rpl10 promotes two RNA conformational changes in the Rpl10-inserted (RI) particle. H38 is retracted away from the eIF5A domain of Nmd3 to adopt its mature position, where it makes extensive contacts with Rpl10 (compare Fig. 1F with 1E, Movie S5). Additionally, there is a subtle (~10 Å) yet important retraction of the middle portion of H89 towards Rpl10, which drives H89 into its mature position to stabilize Rpl10 in its binding cleft between H38 and H89 (Fig. 3, E and F, Movie S6). Unexpectedly, Nmd3 remains in place in progression from the LE to the RI particle (Fig. 1F), suggesting that the insertion of Rpl10, alone, is not sufficient to displace Nmd3 as previously proposed (*17*).

To test the model that Nmd3 holds open RNA helices to promote Rpl10 loading, we mutated residues in Nmd3 that directly contact H38, based on the PL structure (Fig. 4A). We then asked if these mutants could suppress *rpl10-G161D*, a temperature sensitive mutation that destabilizes Rpl10 binding to the ribosome. All structure-guided mutations, including mutation of Arg333, which interacts with the flipped-out base A1025 at the tip of H38, suppressed *rpl10-G161D* to varying degrees (Fig. 4C). Additionally, mutations in Nmd3 that suppress *rpl10-G161D*, which had been identified in early genetic screens (*25, 26*), map to the saddle of the eIF5A-like domain that interacts with H38 and to the histidine thumb of Nmd3 that interacts with H89 in our structure (Fig. 4, A and B). These mutations were highly specific to *rpl10-G161D* as they did not suppress a different mutation in *RPL10*, *rpl10-R98S* that blocks Nmd3 release after Rpl10 insertion (fig. S8) (*27*). Importantly, these suppressing mutations weaken the affinity of Nmd3 to the 60S (*21*), which we can now unambiguously attribute to weakened binding to H38 and H89. Taken together, these results suggest that the release of H38 and H89 from Nmd3 stabilizes Rpl10 in its binding cleft. Thus, the export adapter Nmd3 plays a critical role in both priming the binding site for Rpl10 loading and stabilizing Rpl10 in the ribosome to complete the PTC.

**Fig. 4.**
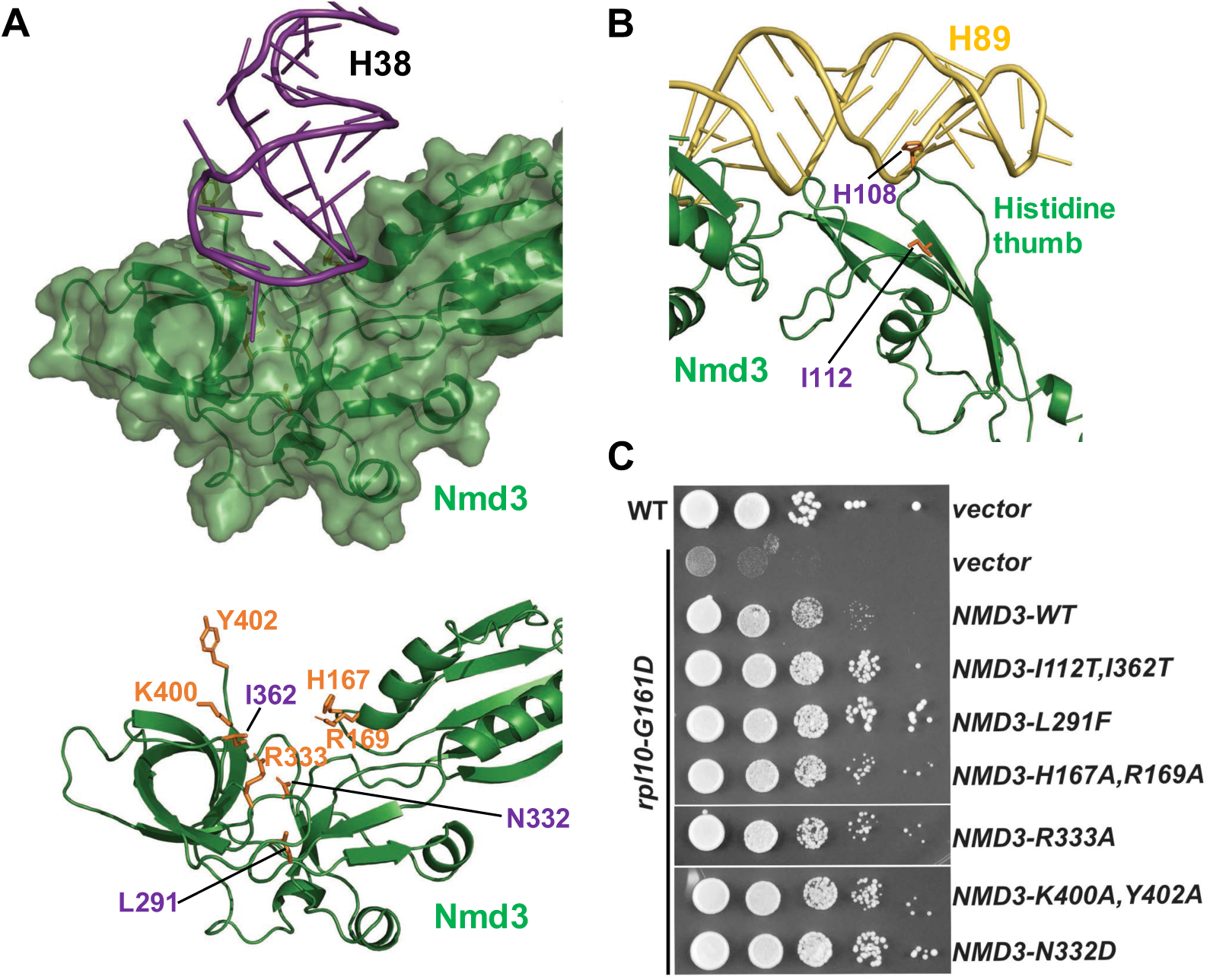
Release of H38 and H89 from Nmd3 stabilizes Rpl10 in the ribosome. **(A)** Atomic structure showing that H38 lays in a saddle of Nmd3 (top). Lower panel, selected residues highlighted in orange sit in the immediate interface between Nmd3 and H38. L291, N332 and I362 (purple) were previously identified from genetic screens for suppressors the temperature-sensitive *rpl10-G161D* mutant. **(B)** Atomic structure showing that interaction of the histidine thumb of Nmd3 with H89. H108 and I112 were identified from genetic screens *rpl10-G161D* suppressors. **(C)** Wild-type or *rpl10-G161D* mutant cells were transformed with empty vector or vector expressing WT or the indicated NMD3 mutants. Ten-fold serial dilutions of cultures were plated onto selective plates and incubated for 2 days at 30°C, a semi-permissive temperature for rpl10-G161D. The mutations H167A, R169A, R333A, K400A, and Y402A in NMD3 were engineered based on the structural information as indicated in (A).

### Conclusions

This study presents the structural framework of cytoplasmic 60S maturation including completion of the PTC and assembly of the P stalk. Importantly, our structures reveal insights into the relative timing of events on the joining face of the subunit, including the exchange of Nog2 for Nmd3 and Nog1 for Yvh1, and the subsequent release of Mrt4 by Yvh1 to promote assembly of the P stalk. We can now propose a comprehensive, detailed model for catalytic center assembly of the ribosome (fig. S9). In the late nuclear particle, loading of the nuclear export adapter Nmd3 is blocked by Nog2 while H89 is displaced by the NTD of Nog1 in the A site. In the earliest pre-60S particle trapped after export (early cytoplasmic-immediate), Nmd3 has closed the L1 stalk and captured H38 while H89 remains displaced by Nog1. In the early cytoplasmic-late particle, the NTD of Nog1 has been released from the A site, allowing the NTD of Nmd3 to dock on Tif6 and bring H89 to its nearly-mature position. In the pre-Lsg1 particle, multiple assembly factors have been released, allowing the loading of Yvh1, which allosterically releases Mrt4 from the P stalk to initiate P stalk assembly. In the Lsg1-engaged particle, the NTD of Nmd3 undergoes a rotation away from H89. This rearrangement releases H89 from Nmd3 to prime the subunit for the insertion of Rpl10. The insertion of Rpl10 causes the retraction of both H38 and H89 to their mature positions, stabilizing Rpl10 in its binding site. The dissociation of H89 and H38 from Nmd3, associated with Lsg1 binding and Rpl10 insertion, respectively, initiates the release of Nmd3 from the pre-60S subunit. After completion of cytoplasmic assembly delineated here and the addition of P proteins to complete the stalk, the nascent 60S subunit undergoes a test drive, which assesses the integrity of the P site and the ability of the 60S subunit to engage and activate GTPases (*11, 12*). Completion of the test drive releases the final biogenesis factors to license the new subunit for *bona fide* translation.

## Acknowledgements

We thank J. Huibregtse and A. Matouschek for critical reading of the manuscript; A. Dai for help with data collection and processing; D. Wrapp, M. Liu, N. Wang, M. Gilman and J. McLellan for support with model building; J. Yelland for help with data collection; and members of the Johnson and Taylor labs for helpful discussions. EM data were acquired at the Sauer Structural Biology Laboratory at UT Austin.

## Author contributions

Y.Z. performed electron microscopy, single particle processing, and model building. S.M. purified the pre-60S samples and performed genetic studies. Y.Z., S.M., A.W.J., and D.W.T. analyzed and interpreted the data and wrote the manuscript. A.W.J. and D.W.T. supervised the study and secured funding for the work.

## Data and materials availability

The cryo-EM structures of LN, ECI, ECL, PL, LE and RI pre-60S intermediates have been deposited into the Electron Microscopy Data Bank with accession numbers EMD-AAAA, EMD-AAAA, EMD-AAAA, EMD-AAAA, EMD-AAAA, and EMD-AAAA, respectively. Their associated atomic models have been deposited into the Protein Data Bank with PDB codes ZZZZ, ZZZZ, ZZZZ, ZZZZ, ZZZZ, and ZZZZ, respectively.

## Supplementary Materials

### Materials and Methods

#### Cell growth

All cells were grown at 30°C in appropriate dropout medium supplemented with 2% glucose or 1% galactose as the carbon source. Site-directed mutations in *NMD3* were generated by inverse PCR in the vector pAJ123. Suppression of *rpl10-G161D* was tested by transforming plasmids into AJY1657 and plating on Leu-deficient media with glucose.

#### Affinity purification of the Rlp24∆C-TAP particles

A cell culture of strain AJY1134 (with mutation reg1-501 that eliminates glucose repression of GAL genes) containing pAJ3965 was grown to OD_600_ of 0.3 in 1.5L of Ura-deficient media with glucose. Galactose was added to a final 1% (w/v) concentration and cells were grown for two more hours, harvested and frozen. Cell pellets were stored at −80°C. Approximately one third of the cell pellet was used per affinity purification. Cell pellets were washed and resuspended in 1.5 volumes of lysis buffer (20mM HEPES, pH 7.5, 10mM MgCl_2_,100mM KCl, 5mM β-mercaptoethanol, 1mM PMSF and protease inhibitors). Extracts were prepared by glass bead lysis and clarified by centrifugation at 4°C for 15 minutes at 18,000g. NP-40 was added to a final concentration of 0.15%(v/v) to the clarified extract which was then incubated with rabbit IgG (Sigma) coupled Dynabeads (Invitrogen) for 1h at 4°C. The Dynabeads were prepared as previously described (*28*). Beads were then washed once with lysis buffer containing 0.15% NP40 and 1 mM dithiothreitol instead of β-mercaptoethanol at 4°C for 5 minutes, and twice with the same buffer but without NP-40. The bound complexes were enzymatically eluted with tobacco etch virus protease for 90 minutes at 16°C. Eluted particles were flash frozen in 10ul aliquots in liquid N2 and then stored at −80°C for cryo-EM.

#### Purification of the Nmd3-TAP particles from diazaborine-treated cells

Cells with C-terminally TAP-tagged Nmd3 (AJY1874) were grown in 1L of YPD media to OD_600_ of 0.4. Diazaborine (Millipore) dissolved in DMSO was added to the cell culture to obtain a final concentration of 10μg/ml and cultures were shaken at 30°C for 30minutes. Diazaborine treated cell cultures were divided into three equal parts and cells were harvested by centrifugation. Cell pellets were stored at −80°C without washing until further use. For purification of diazaborine-treated particles, cell pellets were washed in buffer without diazaborine and particles were purified as described above.

#### Cryo-EM grid preparation and data collection

C-flat holy carbon grids (CF-1.2/1.3, Protochips Inc.) were pre-coated with a thin layer of freshly prepared carbon film and glow-discharged for 30 seconds using a Gatan Solarus plasma cleaner before addition of sample. 2.5 μl of affinity-purified particles (~50 nM for the Rlp24∆C-TAP particles and ~100 nM for the Nmd3-TAP particles isolated from diazaborine-treated cells) were placed onto grids, blotted for 3 seconds with a blotting force of 5 and rapidly plunged into liquid ethane using a FEI Vitrobot MarkIV operated at 4 °C and 100% humidity. Data were acquired using a FEI Titan Krios transmission electron microscope (Sauer Structural Biology Laboratory, University of Texas at Austin) operating at 300 keV at a nominal magnification of ×22,500 (1.1 Å pixel size) with defocus ranging from −1.0 to −2.5 μm. The data were collected using a total exposure of 6 s fractionated into 20 frames (300 ms per frame) with a dose rate of ~8 electrons per pixel per second and a total exposure dose of ~40 e^−^Å^−2^. A total of 7,403 and 6,179 micrographs of the Rlp24∆C-TAP particles and the Nmd3-TAP particles purified from diazaborine-treated cells, respectively, were automatically recorded on a Gatan K2 Summit direct electron detector operated in counting mode using the MSI-Template application within the automated macromolecular microscopy software LEGINON (*29*).

#### Cryo-EM data processing

All image pre-processing was performed in Appion (*30*). Individual movie frames were aligned and averaged using ‘MotionCor2’ drift-correction software (*31*). These drift-corrected micrographs were binned by 8, and bad micrographs and/or regions of micrographs were removed using the ‘manual masking’ command within Appion. A total of 326,567 and 485,448 particles of the Rlp24∆C-TAP particles and the Nmd3-TAP particles from diazaborine-treated cells were picked with a template-based particle picker using a reference-free 2D class average from a small subset of manually picked particles as templates, respectively. The contrast transfer function (CTF) of each micrograph was estimated using CTFFIND4 (*32*). Selected particles were extracted from micrographs using particle extraction within RELION (*33*) and the EMAN2 coordinates exported from Appion.

The workflow for data processing is summarized in figs. S2 and S4. Briefly, between two and three rounds of reference-free 2D classification with 100 classes for each sample were performed in RELION to remove junk particles, resulting in a clean stack of 216,030 particles for the Rlp24∆C-TAP sample purified from yeast cells and 393,665 particles for the Nmd3-TAP sample purified from diazaborine-treated yeast cells. Using a previously published pre-60S structure (EMD-8368) (*18*) low-pass filtered to 60-Å as a reference, 3D classification was performed within RELION to further clean up the particles. The Rlp24∆C-TAP particles were subjected to two rounds of 3D classification and an additional round of 2D classification, resulting in 178,310 ‘good’ particles for further processing. The Nmd3-TAP particles were subjected to one round of 3D classification, resulting in 340,712 ‘good’ particles for the final processing steps. Next, 3D auto-refine was performed in RELION using the best corresponding 3D class model low-pass filtered to 60-Å as a starting model to produce maps at 3.8 Å and 3.7 Å resolution (using the 0.143 gold-standard FSC criterion) for the Rlp24∆C-TAP and the Nmd3-TAP particles, respectively. In order to improve the resolution, movie refinement was also performed using RELION (*34*) with a running average window of 3 movie frames and a standard deviation of 1 pixel for the translational alignment, followed by ‘particle polishing’ (*35*) in RELION to correct for beam-induced motion of individual groups of particles (200 pixel standard deviation) and to perform B-factor weighting using a running average window of 3 frames. These polished particles were then used for another round of 3D auto-refine, resulting in reconstructions both at 3.6 Å resolution (again using the gold-standard FSC from two independent half-maps) for the Rlp24∆C-TAP and the Nmd3-TAP particles. To improve the local density of important regions of the map and to separate various states, a series of focused 3D classification and signal subtraction were performed on the two data sets (Figs. S2 and S5), followed by 3D auto-refine of selected particles. Finally, selected 3D auto-refine maps were subjected to post-processing within RELION with a soft mask and a B-factor of −40 Å^2^. The overall resolution of each structure was calculated based on the gold standard 0.143 FSC criterion using two independent half-maps. The local resolution was also estimated using RELION (figs. S1 and S3). The final reconstructions were segmented using Segger (*36*) in UCSF Chimera (*37*).

#### Atomic model building and refinement

The majority of the model building was performed by manually adjusting and extending existing atomic structures, which had been rigid-body docked into our EM density in Chimera. The atomic structure of a nuclear pre-60S intermediate (PDB 3JCT) (*16*) was used as the initial model for modeling the LN, ECI and ECL structures with Rpl1, Rpl29 and Rpl42 from PDB 5T62 and Rpl12 from PDB 4V88, and part of the rRNAs from PDB 5T62 (*18*) and PDB 4V88 (*38*). Rpl12 is not modeled in the ECI and ECL structures due to poor density. The atomic structure of a cytoplasmic pre-60S particle (PDB 5T62) (*18*) was used as the initial model for modeling the PL, LE and RI structures with Rpl12 and part of the rRNAs from PDB 4V88 (*38*). The structures were optimized by manually inspecting and adjusting the RNA backbones/bases and side chains of amino acids into the corresponding cryo-EM density in Coot (*39*). Yvh1 was built completely de novo by tracing the chain first and then using large side-chains to register the primary sequence within the map. The zinc-binding domains were also used as anchors and sanity checks for our modeling. The final atomic model was refined with real-space refinement (*40*) in PHENIX (*41*) and the refinement statistics are shown in Supplementary Table S1. All structural analysis and figure preparation were performed with UCSF Chimera, Coot and PyMOL (*42*).

## Supplemental Movie Legends

**Movie S1. Architecture of 60S precursors.** This movie shows each of the six atomic models displayed in Fig.1 rocked by 30-degrees around the y-axis twice. The Rpl12 structure is docked into ECI and ECL models.

**Movie S2. Rearrangements of ribosomal proteins and RNA during maturation of the 60S.** This movie is an animation morphing between the structures in Fig 1. The sequence is the same as in Movie S1. The Rpl12 structure is docked into ECI and ECL models.

**Movie S3. Large-scale rearrangement of H89.** This movie shows the large-scale rearrangement of H89 to its nearly-mature position where it engages with the histidine thumb of Nmd3 upon the release of the N-terminus of Nog1 from the A site.

**Movie S4. Rotation of the NTD of Nmd3.** This movie shows the 60-degree rotation of the NTD of Nmd3 upon Lsg1 engagement by interpolation between Nmd3 structures from the PL particle and the LE particle. The eL22 and eIF5A domains of Nmd3 remain relatively rigid.

**Movie S5. Retraction of H38 by Rpl10.** This movie shows that retraction of H38 away from the eIF5A domain of Nmd3 to adopt its mature position upon Rpl10 insertion.

**Movie S6. Retraction of H89 by Rpl10.** This movie shows the subtle (~10 Å) yet important retraction of the middle portion of H89 towards Rpl10, which drives H89 into its mature position to stabilize Rpl10 in its binding cleft between H38 and H89.

**Table S1.** Statistics of cryo-EM structure determination and model refinement.

| Data collection and processing |  |  |  |  |  |  |
| --- | --- | --- | --- | --- | --- | --- |
| Sample | Rlp24ΔC-TAP pre-60S |  |  | Nmd3-TAP pre-60S from diazaborine-treated cells |  |  |
| Microscope | FEI Titan Krios |  |  | FEI Titan Krios |  |  |
| Direct electron detector | Gatan K2 Summit |  |  | Gatan K2 Summit |  |  |
| Voltage (keV) | 300 |  |  | 300 |  |  |
| Pixel size (Å) | 1.1 |  |  | 1.1 |  |  |
| Total dose (e<sup>-</sup>/ Å<sup>2</sup>) | 40 |  |  | 40 |  |  |
| Defocus range (μm) | 1.0 - 2.5 |  |  | 1.0 - 2.5 |  |  |
| Total micrographs | 7,403 |  |  | 6,179 |  |  |
| Used micrographs | 6,860 |  |  | 5,839 |  |  |
| Total particles | 326,527 |  |  | 485,448 |  |  |
| Particles for initial 3D refinement | 178,310 |  |  | 340,712 |  |  |
| Model building and refinement |  |  |  |  |  |  |
| Model name | LN | ECI | ECL | PL | LE | RI |
| Map resolution at FSC 0.143 (Å) | 3.5 | 3.6 | 3.6 | 3.5 | 3.8 | 3.5 |
| Initial model used (PDB ID) | 3JCT | 3JCT | 3JCT | 5T62 | 5T62 | 5T62 |
| Model composition |  |  |  |  |  |  |
| Peptide chains | 46 | 45 | 45 | 44 | 45 | 46 |
| Residues | 7,767 | 7,441 | 7,094 | 7,010 | 7,387 | 7,594 |
| RNA chains | 3 | 3 | 3 | 3 | 3 | 3 |
| RNA bases | 3,326 | 3,479 | 3,479 | 3,482 | 3,482 | 3,480 |
| R.m.s deviation |  |  |  |  |  |  |
| Bond length (Å) | 0.007 | 0.006 | 0.007 | 0.006 | 0.006 | 0.007 |
| Bond angle (°) | 0.880 | 0.866 | 0.879 | 0.814 | 0.862 | 0.848 |
| Ramachandran plot |  |  |  |  |  |  |
| Outliers (%) | 0.33 | 0.42 | 0.46 | 0.38 | 0.50 | 0.40 |
| Allowed (%) | 7.30 | 6.58 | 7.32 | 6.54 | 6.78 | 7.28 |
| Favored (%) | 92.37 | 93.00 | 92.22 | 93.08 | 92.72 | 92.32 |
| Validation (protein) |  |  |  |  |  |  |
| Clashscore, all atoms | 4.00 | 4.32 | 4.17 | 3.41 | 3.92 | 3.28 |
| Molprobability score | 1.66 | 1.66 | 1.68 | 1.58 | 1.64 | 1.59 |
| Rotamers outlier (%) | 0.51 | 0.51 | 0.64 | 0.43 | 0.48 | 0.56 |
| Validation (RNA) |  |  |  |  |  |  |
| Correct sugar puckers (%) | 98.61 | 98.82 | 98.82 | 99.05 | 98.88 | 99.02 |
| Good backbone conformations (%) | 77.42 | 76.97 | 77.98 | 79.00 | 78.40 | 80.26 |

**Table S2.**
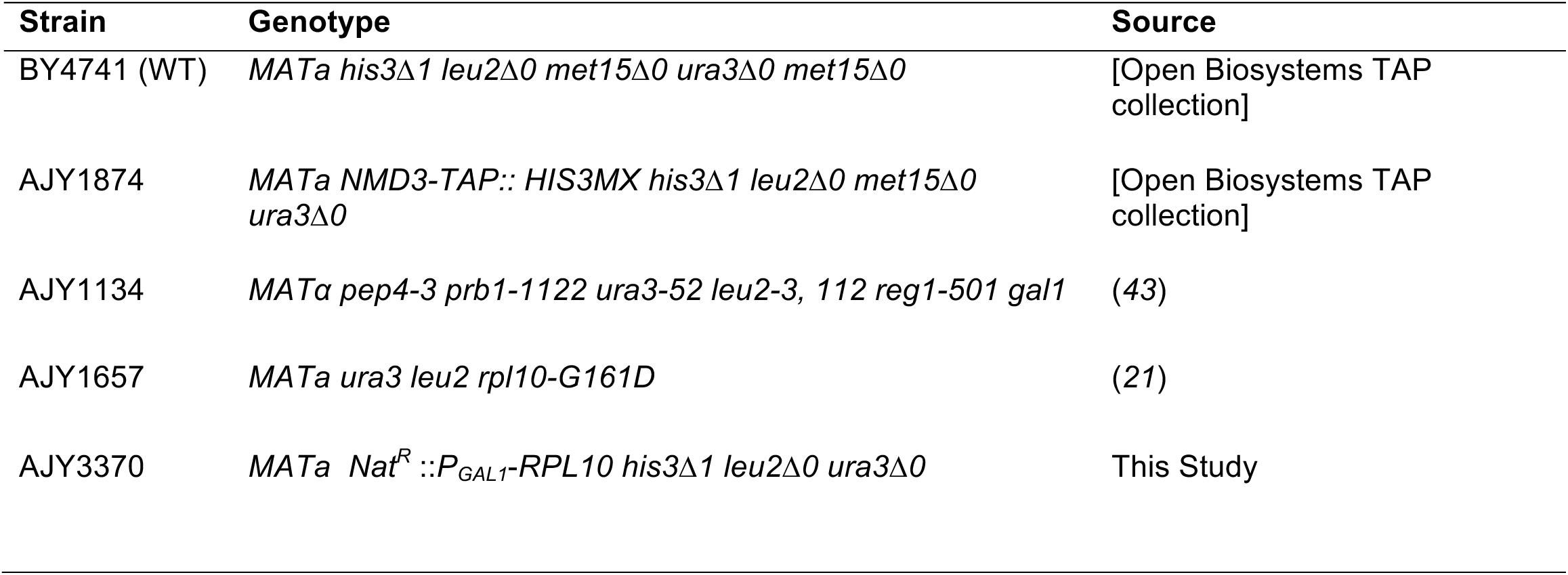
Strains used in this study.

**Table S3.** Plasmids used in this study.

| **Plasmid** | **Description** | **Source** |
| --- | --- | --- |
| pAJ123 | *NMD3 LEU2 CEN* | (44) |
| pAJ415 | *NMD3-L291F LEU2 CEN* | (21) |
| pAJ1334 | *NMD3-I112T, I362T LEU2 CEN* | (21) |
| pAJ2726 | *rpl10-R98S LEU2 CEN* | (27) |
| pAJ3020 | *rpl10-R98S URA3 CEN* | (27) |
| pAJ3609 | *NMD3-C35G URA3 CEN* | (27) |
| pAJ4305 | *NMD3-H167A, R169A LEU2 CEN* | This Study |
| pAJ4307 | *NMD3-R333A LEU2 CEN* | This Study |
| pAJ4308 | *NMD3-K400A, Y402A LEU2 CEN* | This Study |
| pAJ4309 | *NMD3-N332D LEU2 CEN* | This Study |
| pAJ3965 | *GAL10::RPLP24-ΔC-TAP URA3 CEN* | This Study |
| pRS415 | *LEU2 CEN* | (45) |
| pRS416 | *URA3 CEN* | (45) |

**Fig. S1.**
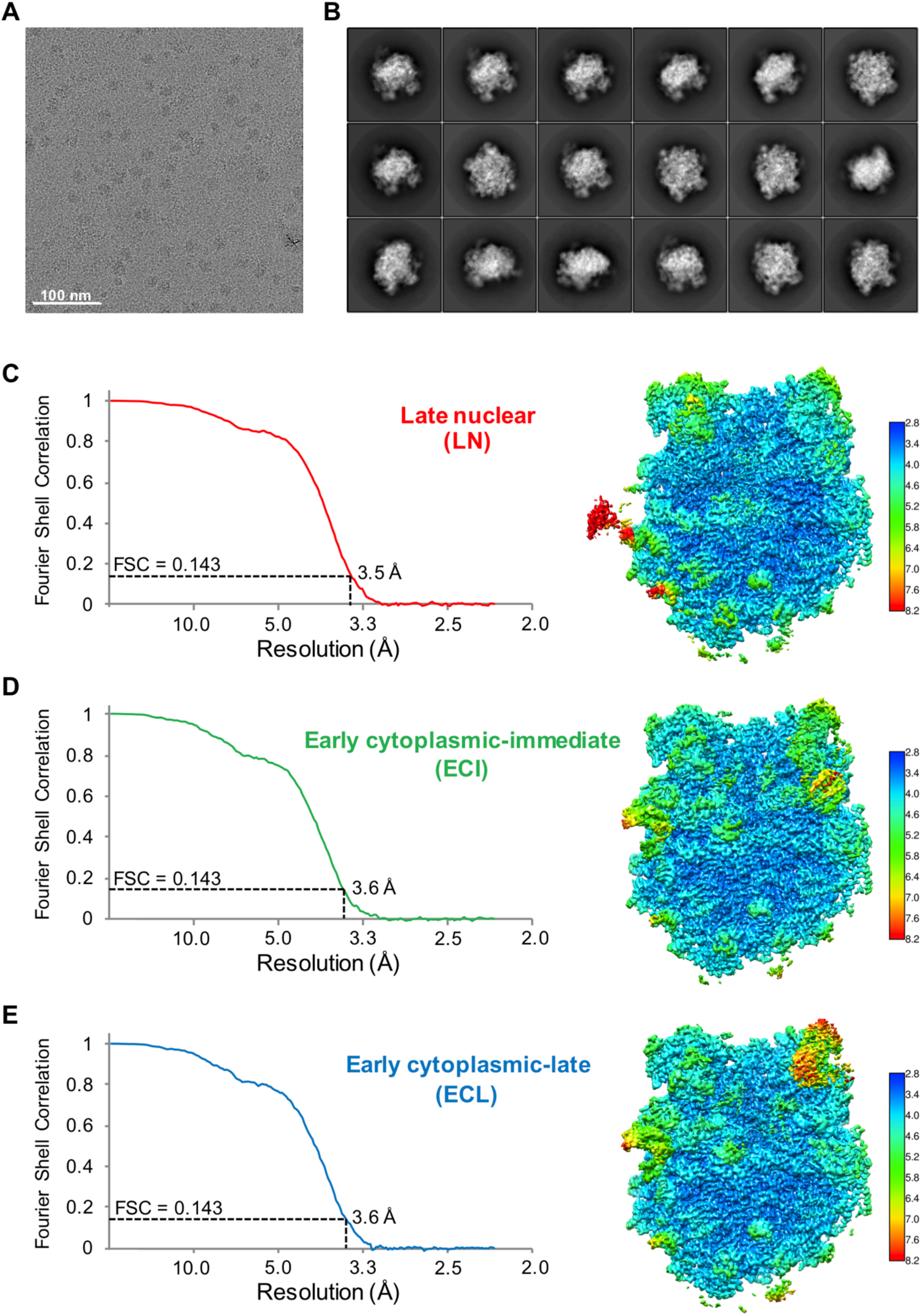
Cryo-EM structure determination of the Rlp24∆C-TAP pre-60S ribosomes. **(A)** A drift-corrected cryo-EM micrograph of the Rlp24∆C-TAP pre-60S ribosomes imaged on a Gatan K2 Summit direct electron detector. The scale bar indicates 100 nm. **(B)** Reference-free 2D class averages of the Rlp24∆C-TAP pre-60S particles. The width of the boxes is 422.4 Å. **(C)** Fourier shell correlation curve (left) and local resolution map (right) for the late nuclear (LN) pre-60S particles. The overall resolution for the map is 3.5-Å based on the 0.143 FSC criterion. **(D)** Fourier shell correlation curve (left) and local resolution map (right) for the early cytoplasmic-immediate (ECI) pre-60S particles. The overall resolution for the map is 3.6-Å based on the 0.143 FSC criterion. **(E)** Fourier shell correlation curve (left) and local resolution map (right) for the early cytoplasmic-late (ECL) pre-60S particles. The overall resolution for the map is 3.6-Å based on the 0.143 FSC criterion. The release of the NTD of Nog1 from the A site likely causes high flexibility of the P stalk, leading to badly resolved Rpl12 and Mrt4 on the P stalk.

**Fig. S2.**
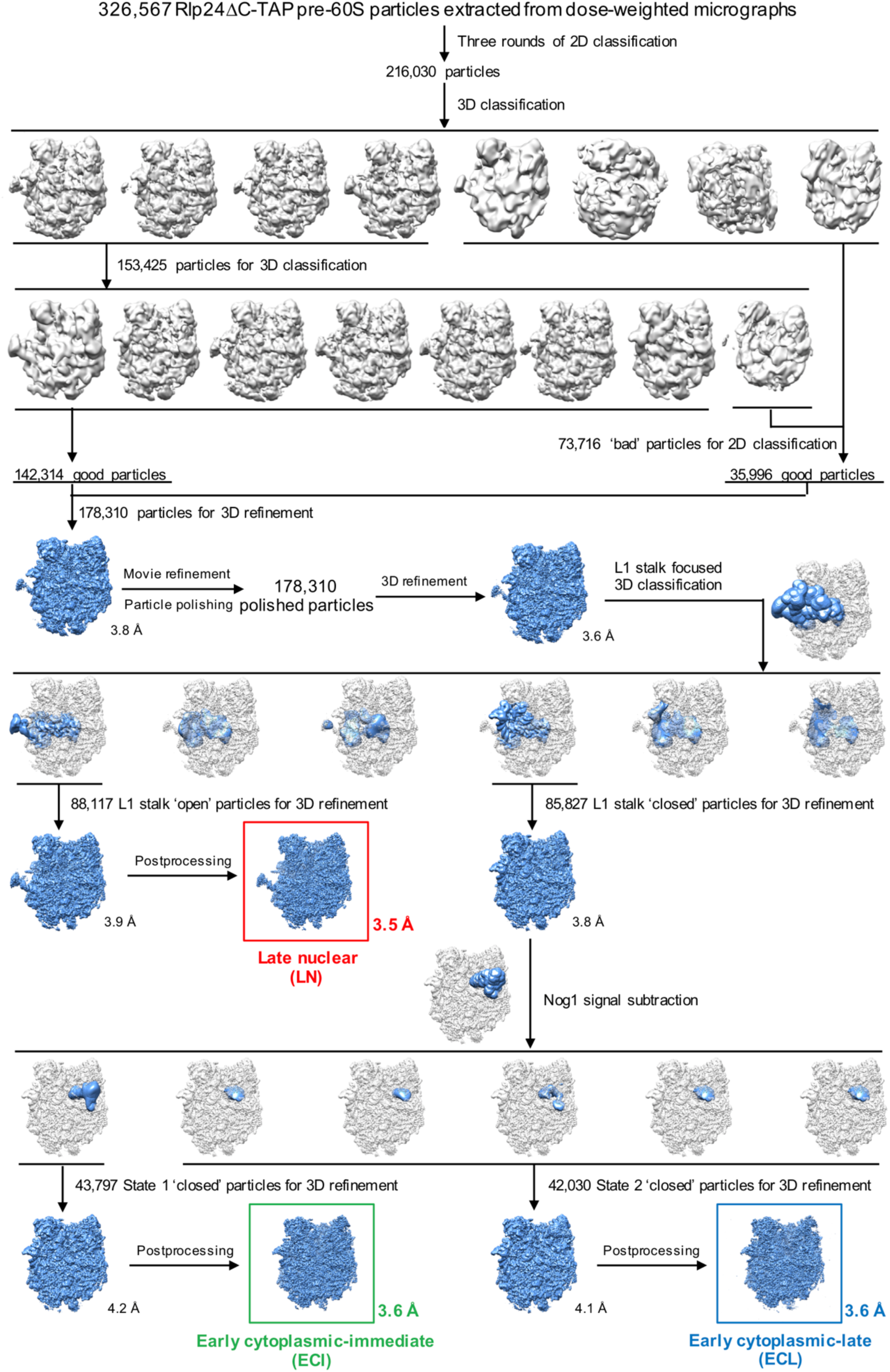
Classification and refinement workflow of the Rlp24∆C-TAP pre-60S ribosomes. A total starting stack of 326,567 Rlp24∆C-TAP particles extracted from dose-weighted micrographs were subjected to three rounds of reference-free 2D classification. A subset of 216,030 particles from the final 2D classes were selected and subjected to two rounds of 3D classification and an additional round of 2D classification, resulting in 178,310 ‘good’ particles which were further subjected to 3D refinement, movie refinement and particle polishing in RELION. Next, the polished particles were subjected to L1 stalk focused 3D classification using a soft mask around the L1 stalk, resulting in 88,117 L1 stalk ‘open’ particles and 85,827 L1 stalk ‘closed’ particles. The L1 stalk ‘open’ particles, which were termed late nuclear (LN) particle in this paper, were further analyzed in RELION, leading to a 3.9-Å map after 3D refinement and a 3.5-Å map after postprocessing. The L1 stalk ‘closed’ particles, which were termed early cytoplasmic (EC) particles in this paper, were further subjected to signal subtraction analysis using a soft mask around the Nog1 N-terminal four-helical-bundle domain and GTPase domain (NTD-GD), resulting in two subsets of particles: 43,797 particles with clear density of NTD-GD (early cytoplasmic-immediate (ECI)) and 42,030 particles with no density of NTD-GD (cytoplasmic-late (ECL)). Further analysis of the ECI particles led to a 4.2-Å 3D refinement map and a 3.6-Å postprocess map, where the NTD-GD of Nog1 is in its canonical position with the N-terminus of Nmd3 being unresolved. In contrast, further analysis of the ECL particles led to a 4.1-Å 3D refinement map and a 3.6-Å postprocess map, where Nog1 NTD-GD has been released from the A site while the N-terminus of Nmd3 has docked onto Tif6.

**Fig. S3.**
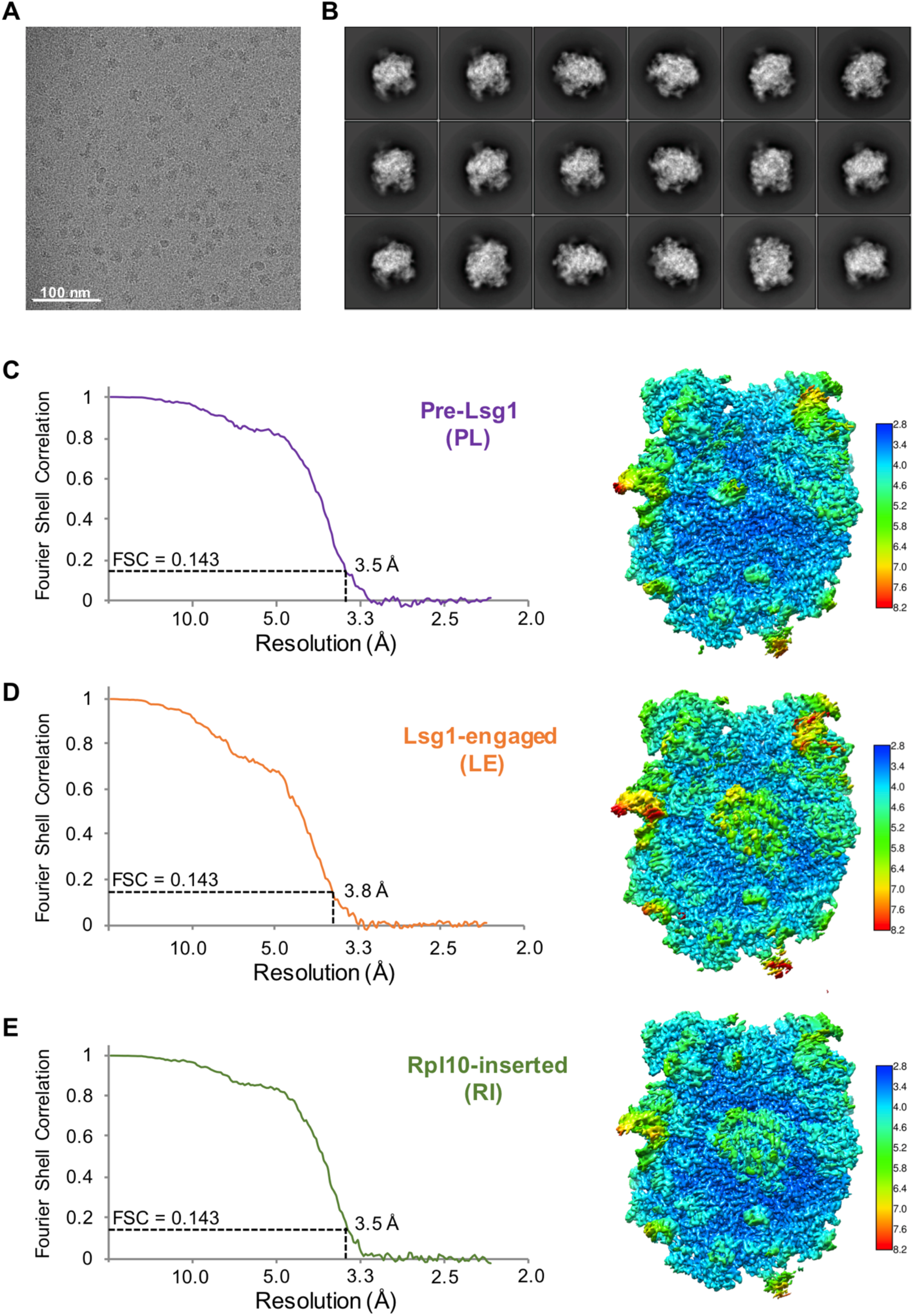
Cryo-EM structure determination of the Nmd3-TAP pre-60S ribosomes from diazaborine-treated cells. **(A)** A drift-corrected cryo-EM micrograph of the Nmd3-TAP pre-60S ribosomes imaged on a Gatan K2 Summit direct electron detector. The scale bar indicates 100 nm. **(B)** Reference-free 2D class averages of the Nmd3-TAP pre-60S particles. The width of the boxes is 422.4 Å. **(C)** Fourier shell correlation curve (left) and local resolution map (right) for the pre-Lsg1 (PL) pre-60S particles. The overall resolution for the map is 3.5-Å based on the 0.143 FSC criterion. **(D)** Fourier shell correlation curve (left) and local resolution map (right) for the Lsg1-engaged (LE) pre-60S particles. The overall resolution for the map is 3.8-Å based on the 0.143 FSC criterion. **(E)** Fourier shell correlation curve (left) and local resolution map (right) for the Rpl10 inserted (RI) pre-60S particles. The overall resolution for the map is 3.5-Å based on the 0.143 FSC criterion.

**Fig. S4.**
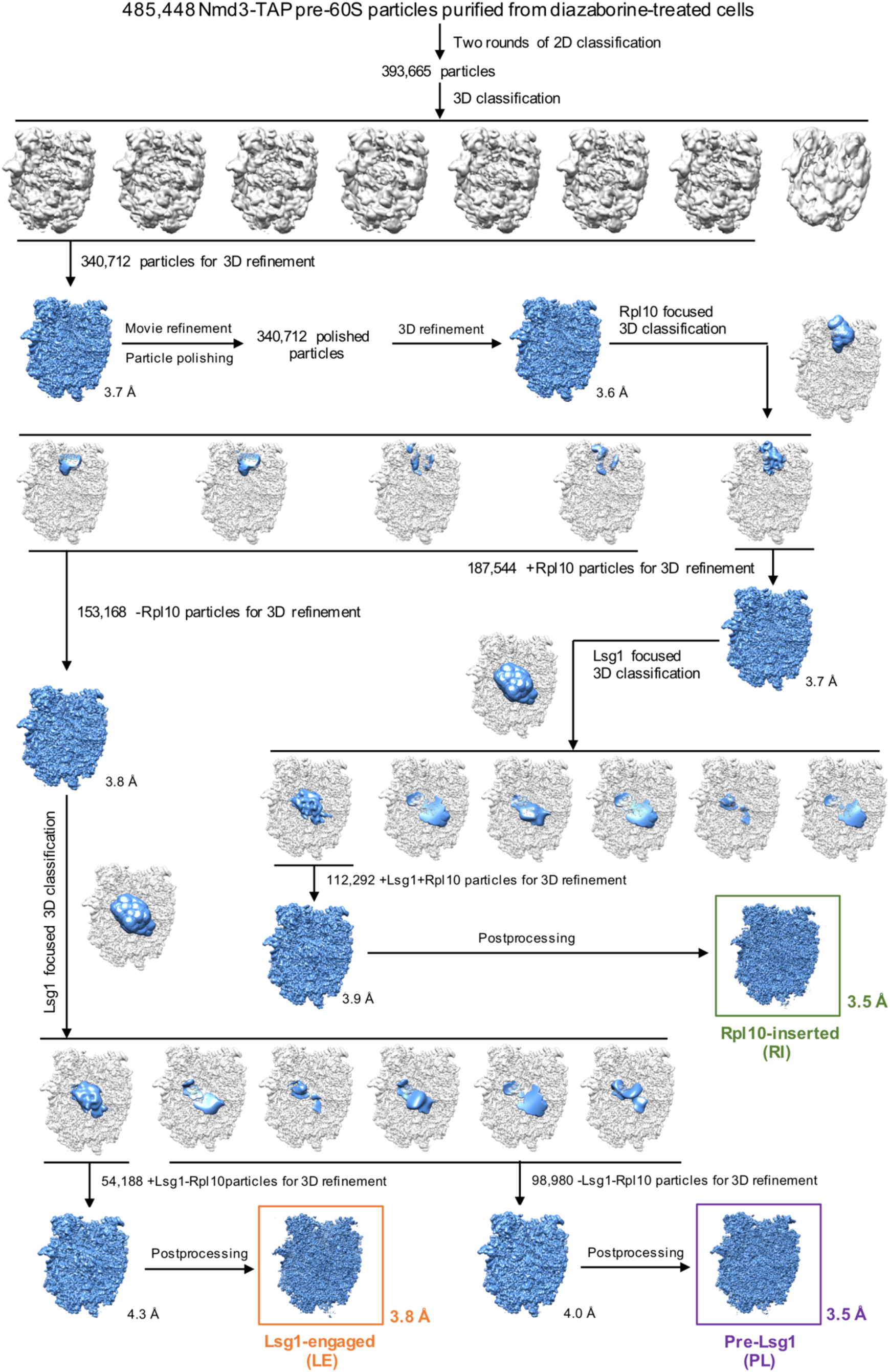
Classification and refinement workflow of the Nmd3-TAP pre-60S ribosomes purified from diazaborine-treated cells. A total starting stack of 485,448 particles were subjected to two rounds of reference-free 2D classification. A subset of 393,665 particles from the final 2D classes were selected and subjected to 3D classification, resulting in 340,712 ‘good’ particles which were further subjected to 3D refinement, movie refinement and particle polishing in RELION. The polished particles were first subjected to focused 3D classification with a soft mask around Rpl10, resulting in 153,168 particles without Rpl10 (-Rpl10) and187,544 particles with Rpl10 (+Rpl10). The-Rpl10 particles were further subjected to Lsg1 focused 3D classification, resulting in 98,980 -Lsg1-Rpl10 particles and 54,188 +Lsg1-Rpl10 particles, which were termed pre-Lsg1 (PL) and Lsg1-engaged (LE) particles, respectively. Further processing of the PL particles led to a 4.0-Å 3D refinement map and a 3.5-Å postprocess map, while further processing of the LE particles resulted in a 4.3-Å 3D refinement map and a 3.8-Å postprocess map. The +Rpl10 particles were also subjected to Lsg1 focused 3D classification, resulting in 112,292 particles with strong Lsg1 density (+Lsg1+Rpl10), which were named Rpl10-inserted (RI) particles. Further processing of the RI particles led to a 3.9-Å 3D refinement map and a 3.5-Å postprocess map.

**Fig. S5.**
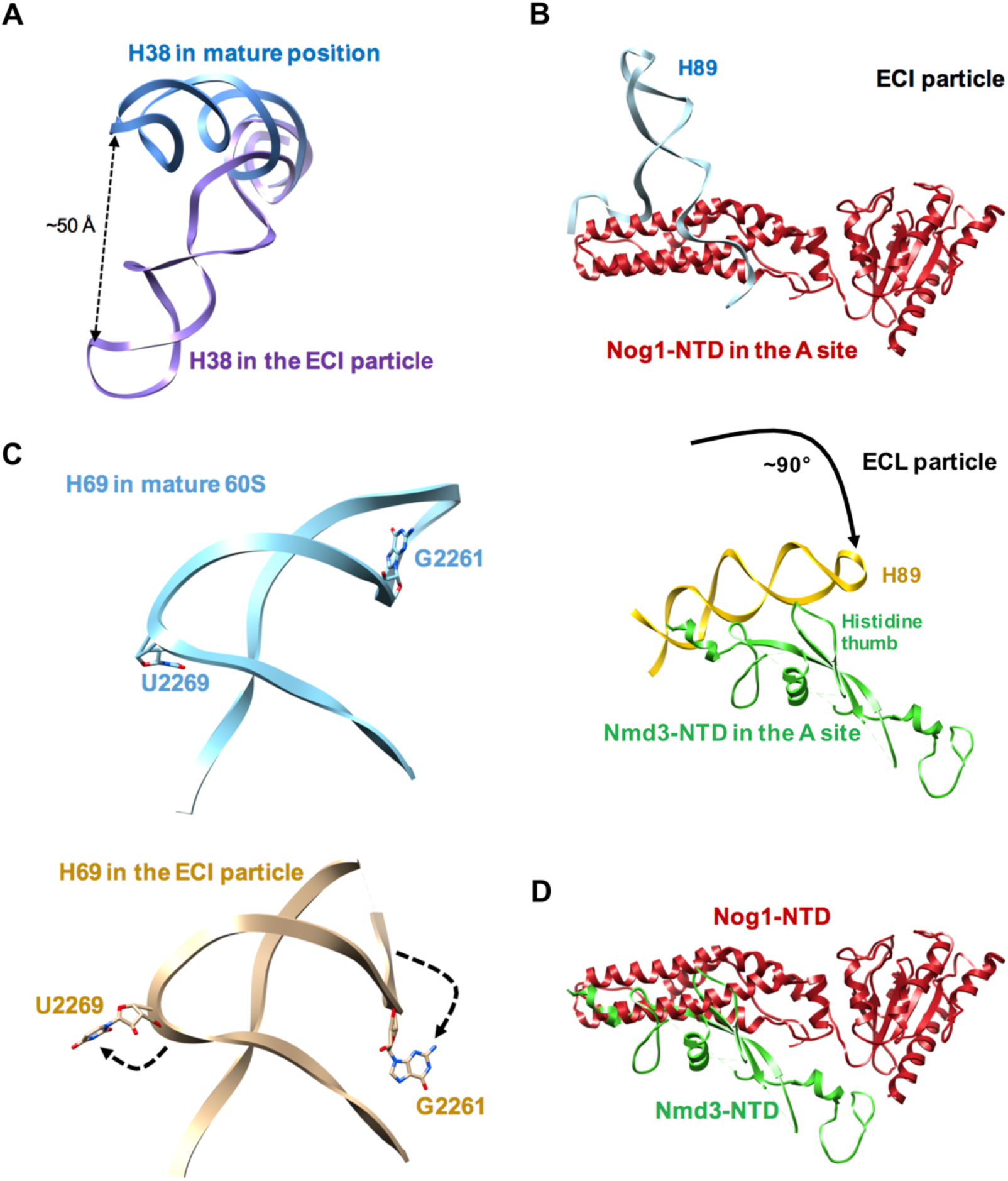
Dynamics of H38, H89 and H69 in pre-60S ribosomes. **(A)** Comparison of H38 in its mature position (PDB: 4V88) (*38*) and H38 from the early cytoplasmic-immediate (ECI) particle. There is a ~50 Å shift of the H38 tip. **(B)** Rearrangement of H89 in transition from ECI (top) to ECL (bottom). Upon the release of Nog1 NTD from the A site and Nmd3 NTD docking onto Tif6, H89 is rearranged to its near-mature position by rotating about ~90° to interact with the histidine thumb of Nmd3. **(C)** Comparison of H69 in mature 60S (top) (PDB: 4V88) (*38*) and H69 from the early cytoplasmic-immediate (ECI) particle (bottom). G2261 and U2269 are flipped out in the ECI particle relative to mature 60S. The extreme tip of H89 in the ECI particle is not modeled due to bad density. **(D)** Structural conflict of Nog1 and Nmd3 on pre-60S particles. Nog1 NTD from the ECI particle and Nmd3 NTD from the early cytoplasmic-late (ECL) particle are overlaid.

**Fig. S6.**
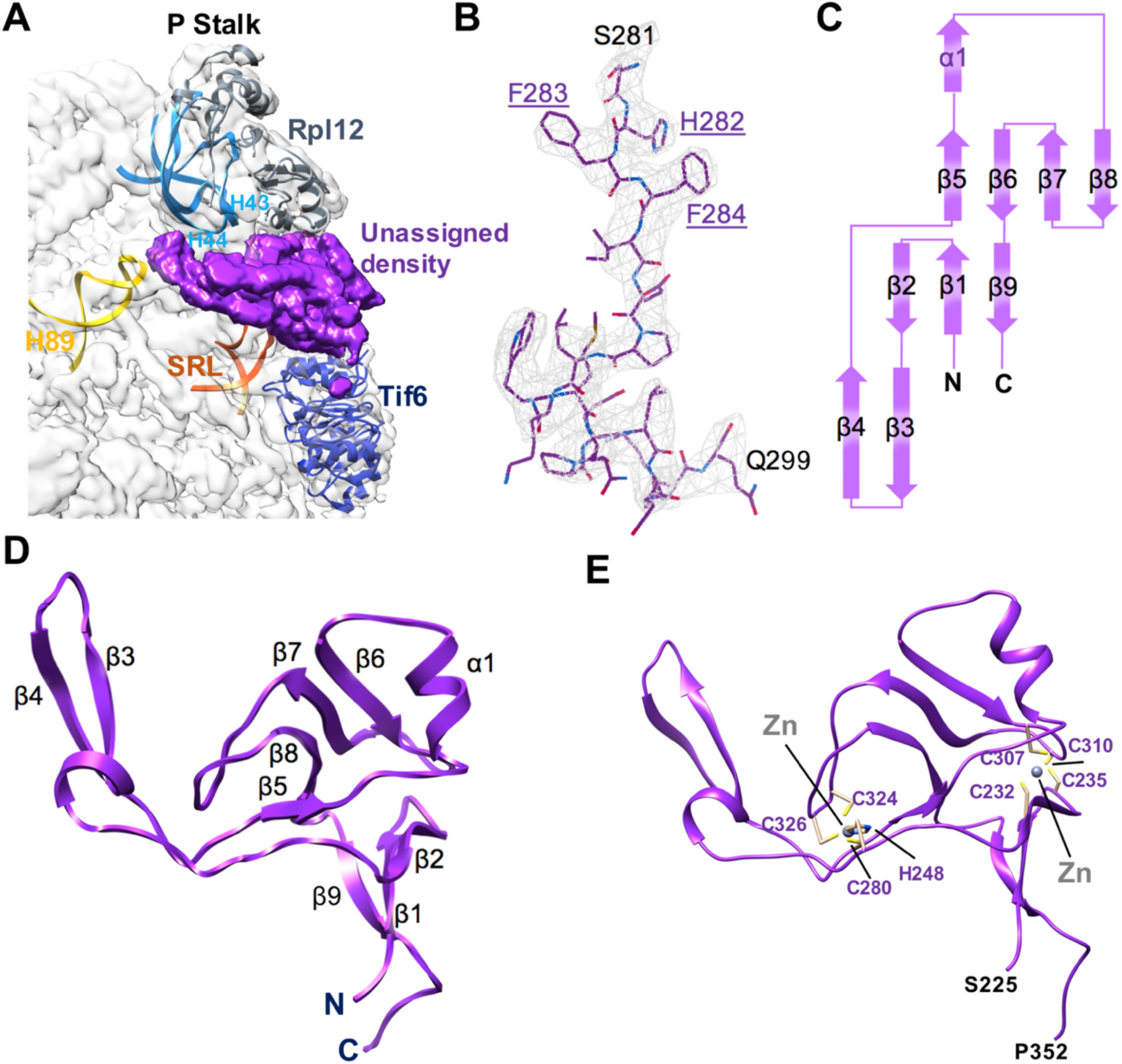
The atomic structure of the zinc-binding domain of Yvh1. **(A)** Segmented unassigned density (purple) positioned between Tif6 and the P stalk in the electron density map of the pre-Lsg1 particle with relevant elements of the 60S subunit indicated. SRL, sarcin-ricin loop. **(B)** Example of the quality of the electron density map showing fitting of side chains of Yvh1 from residues S281 to Q299. Residues H282, F283 and F284 were used as our starting register for model building. **(C)** Topology of the zinc-binding domain of Yvh1. **(D)** Atomic structure of the zinc-binding domain of Yvh1. Secondary structure elements are indicated. **(E)** Detail of the two zinc centers of Yvh1 with coordinating residues indicated and zinc ions modeled in their predicted locations.

**Fig. S7.**
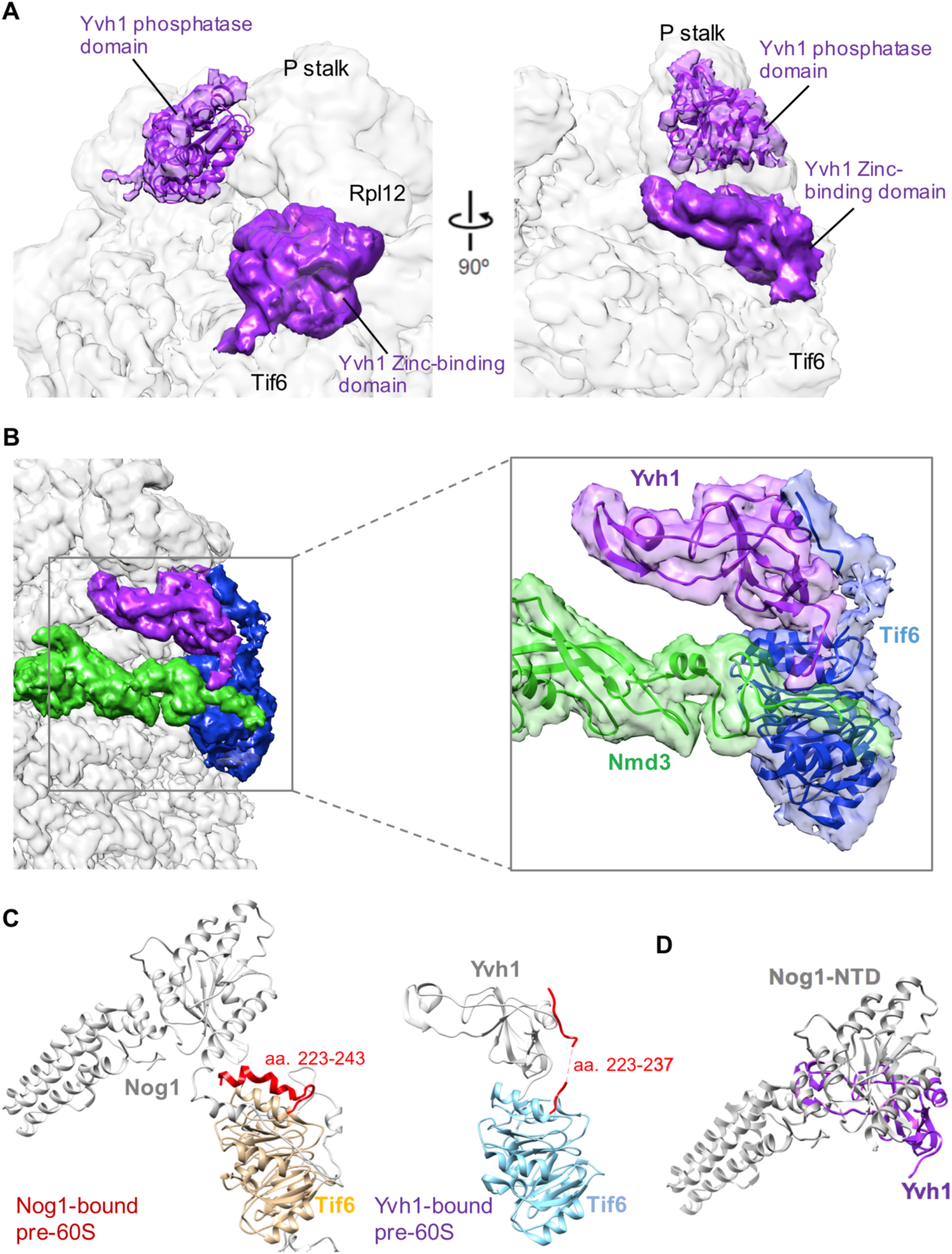
Interactions among Tif6, Nog1, Nmd3 and Yvh1. **(A)** The structure of the Yvh1 phosphatase domain from *Chaetomium thermophilum* (PDB: 5M43) was docked into a density observed at low threshold on the face of the P stalk facing the central protuberance. This roughly corresponds to the previously reported position of the phosphatase domain of Yvh1 (*46*). **(B)** Segmented densities for Yvh1, Nmd3 and Tif6 (left panel) with atomic structures modeled into the densities (right panel). At low threshold, the C-terminus of Tif6 can be traced as a continuous density extending to the surface of Yvh1, although it is hard to model the middle of Tif6 C-terminus. The extreme N-terminus of Nmd3 extends over Tif6 and is crossed by the C-terminus of Yvh1. **(C)** Comparison of the position of the C-terminal tail of Tif6 (aa223-246) in the presence of Nog1 versus Yvh1. The tail folds back over the globular domain of Tif6 in the presence of Nog1 but repositions by rotating ~60° to extend to the surface of Yvh1 in the Yvh1-containing particles. **(D)** Structural conflict of Nog1 and Yvh1 on pre-60S particles. Nog1 from the ECI particle and Yvh1 from the PL particle are overlaid.

**Fig. S8.**
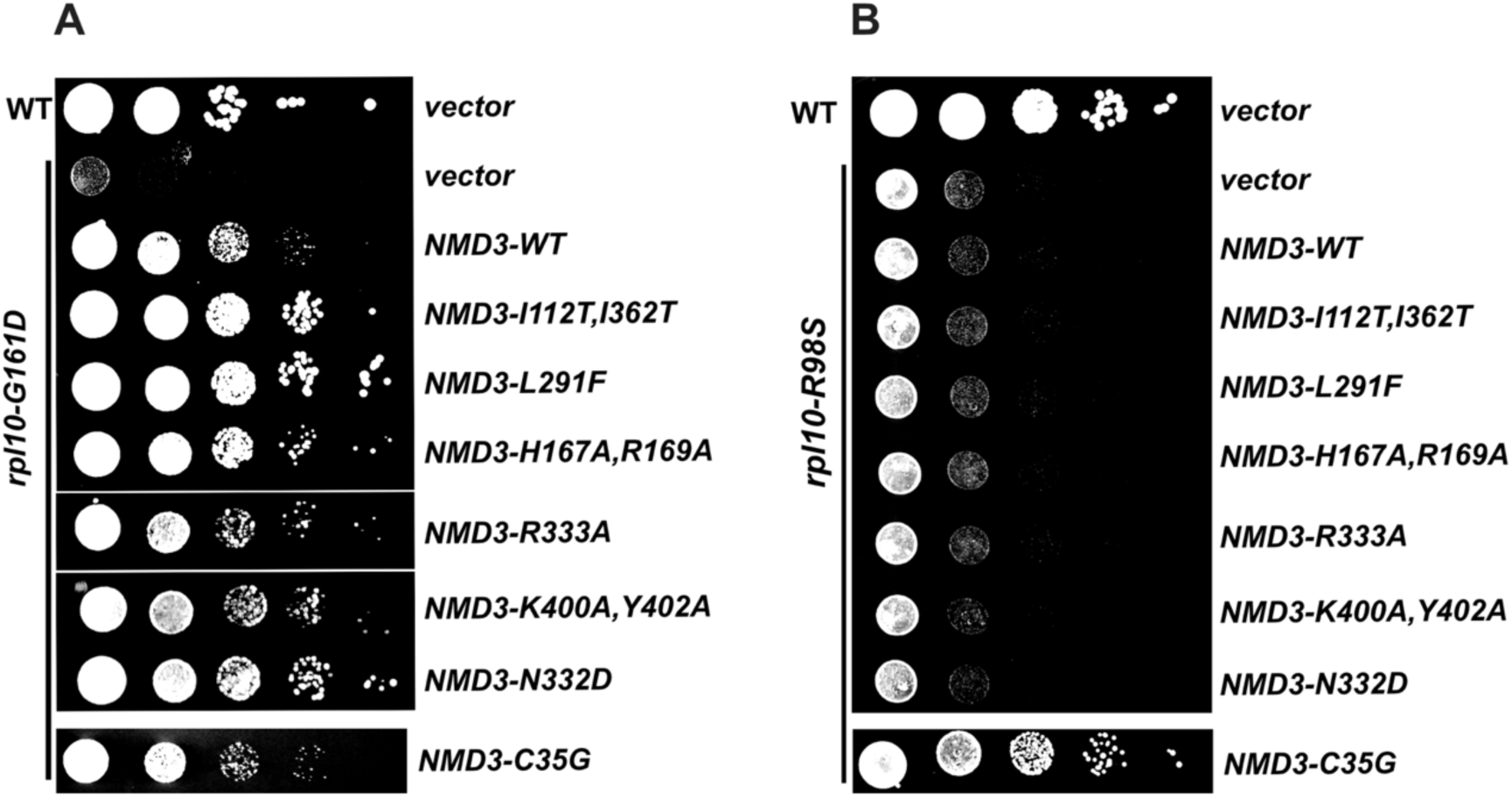
Allele-specific suppression of *rpl10* mutants. **(A)** Wild-type or *rpl10-G161D* mutant cells were transformed with empty vector or vectors expressing WT or mutant Nmd3, as indicated. Ten-fold serial dilutions of cultures were plated onto selective plates and incubated for 2 days at 30°C, a semi-permissive temperature *for rpl10-G161D*. Upper panel is reproduced from Fig. 4C for comparison. **(B)** Wild-type cells were transformed with empty vector and *P_GAL1_-RPL10* mutant cells were co-transformed with an *rpl10-R98S* expressing vector and vectors expressing WT or mutant Nmd3, as indicated. Ten-fold serial dilutions of cultures were plated onto selective plates and incubated for 2 days at 30°C, a semi-permissive temperature for *rpl10-G161D*.

**Fig. S9.**
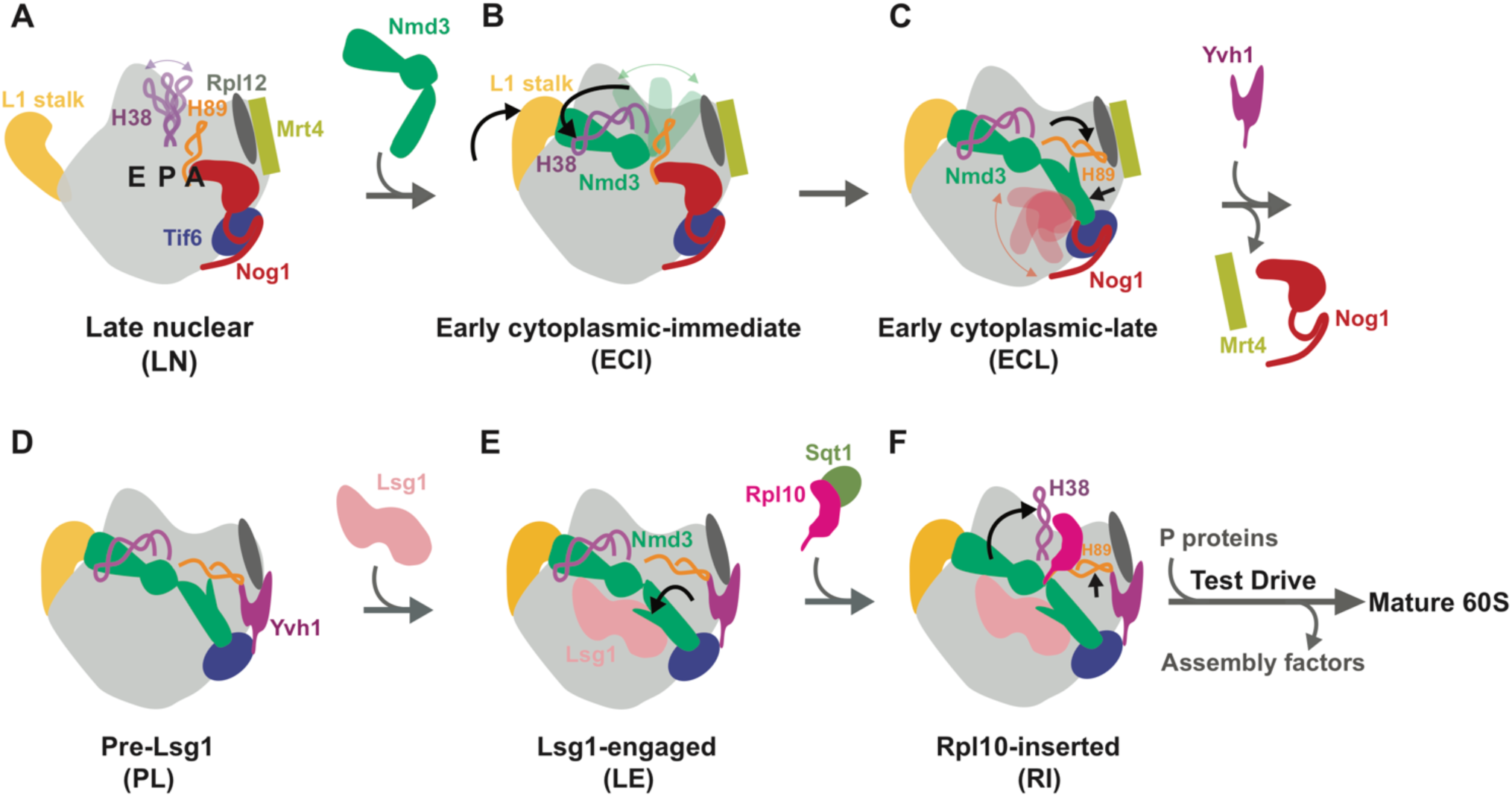
Summary of 60S maturation concluding with Rpl10 insertion. **(A)** Late nuclear particle before Nmd3 loading. H38 tip is flexible while H89 is displaced by the NTD of Nog1 in the A site. Rpl12 and Mrt4 both bind to the P stalk. **(B)** Early cytoplasmic-immediate particle showing that the nuclear export factor Nmd3 closes the L1 stalk and captures H38. The NTD of Nog1 remains in the A site, displacing H89, and the NTD of Nmd3 is not discernible, probably because of its high mobility before docking onto Tif6. **(C)** In the early cytoplasmic-late particle, the NTD and GTPase domain of Nog1 have been released from the A site, allowing Nmd3 to dock on Tif6 and bind H89, which is rearranged to its near-mature position. Note that the Nog1 C-terminus remains in place while the NTD and GTPase domain are not discernible, likely due to high flexibility after displacement from the A site. **(D)** In the Pre-Lsg1 particle, multiple assembly factors have been released. The release of Nog1 allows the binding of Yvh1 to release Mrt4 from the P stalk. **(E)** The Lsg1-engaged particle reveals a rotation of the NTD of Nmd3 away from H89 to engage with Lsg1. This releases H89 from Nmd3 to prime the subunit for the insertion of Rpl10. **(F)** The insertion of Rpl10 causes retraction of both H38 and H89 to their mature positions to stabilize Rpl10 in its binding site. The complete release of Nmd3 from H38 and H89 poises Nmd3 for its imminent release. Subsequently, the nascent 60S subunit undergoes a test drive using molecular mimics of translation factors before licensing it for *bona fide* translation.

